# A largely shared genetic architecture underlies *Saccharomyces cerevisiae* persistence across different mouse host genotypes

**DOI:** 10.64898/2026.11.04.764303

**Authors:** Brandon Bernardo, Yunsun Eoh, Matthew D. Dean, Ian M. Ehrenreich

## Abstract

The extent to which interactions between microbial and host genotypes shape opportunistic pathogen persistence within hosts is an important open question. To assess the contribution of these interactions to persistence, we used *Saccharomyces cerevisiae*, a genetically tractable model for studying opportunistic fungal persistence in mammalian hosts. A pooled panel of barcoded haploid segregants from a cross between a laboratory strain and a clinical isolate, together with both parents, was introduced intravenously into seven genetically diverse mouse strains. We measured the relative persistence of segregants in mouse brains and livers using barcode sequencing. Persistence showed transgressive segregation in both organs. On average across the two organs, yeast genotype explained 45.2% of the variance in segregant persistence, whereas yeast genotype-by-host genotype interactions explained only 2.9%. Linkage mapping showed that loci detected in more host genotypes generally explained more phenotypic variance than those detected in fewer host genotypes. At some loci, genetic variation correlated with increased yeast persistence across multiple organs (general loci) while at others increased persistence in one organ came at the cost of decreased persistence in another (antagonistic loci). General loci contributed disproportionately to brain persistence, whereas liver persistence reflected contributions from both classes. The clinical parent showed greater persistence than the laboratory parent in every host-organ combination, and the additive effects of detected loci predicted the direction of this advantage. Overall, this work shows that in this system, relative microbial persistence is polygenic, with a strong additive component and only a very small contribution from microbial genotype-by-host genotype interactions.

**Article summary:** Whether loci that enable fungi to persist in one host have similar effects in other genetically distinct hosts is unclear. To address this, we introduced a pool of hundreds of genetically distinct yeast segregants and their parent strains into seven genetically diverse mouse hosts. We measured how well each yeast strain persisted in the brain and liver of each mouse. Differences in persistence between yeast strains in the brain and liver were observed across all hosts. The same loci explained organ-specific persistence across hosts, and yeast strains that persisted well in one mouse strain generally did so in the others.

## Introduction

Fungi are a major class of opportunistic human pathogens that cause substantial morbidity and mortality, particularly in immunocompromised individuals (Brown et al. 2012; Köhler et al. 2015; Almeida et al. 2019). Because these infections are often difficult to treat, identifying mechanisms that enable fungi to persist within mammalian hosts is important for understanding pathogenesis and developing more effective therapies (Scorzoni et al. 2017; Ciurea et al. 2020; Nivoix et al. 2020; van de Veerdonk et al. 2025). Fungal persistence within a host can depend on genetic variation in both the fungus and its host, as well as interactions between their genotypes (Byron et al. 1995; Marakalala et al. 2013). However, the relative contributions of fungal genotype and fungal genotype-by-host genotype interactions to persistence remain poorly understood. Such interactions could limit how broadly genetic effects identified in one host background apply to others. Clarifying these contributions is therefore important for identifying general mechanisms of persistence and understanding variation in infection outcomes.

The budding yeast *Saccharomyces cerevisiae* offers a tractable system for understanding how fungal genetic variation shapes persistence across mammalian host genotypes (Goldstein and McCusker 2001). Although generally benign and best known as a model organism and agent of fermentation, *S. cerevisiae* can cause opportunistic infections (Strope et al. 2015). Clinical isolates of *S. cerevisiae* often show the ability to survive and proliferate in mammalian hosts (McCusker 2006). Studies in mice have shown that pathogenic potential differs among isolates and have connected this variation to traits including growth at elevated temperatures and pseudohyphal development (Clemons et al. 1994; McCusker et al. 1994). A recent comparative genomic study further found that infective isolates were distributed among multiple lineages, including populations associated with food production and probiotic use (Morard et al. 2023). *S. cerevisiae* is also highly amenable to genetic analysis. Crosses can generate large panels of haploid meiotic progeny with known genotypes, and chromosomally integrated DNA barcodes allow the relative abundance of hundreds of genotypes to be measured simultaneously within a host (Matsui et al. 2022; Mullis et al. 2022). Together, these features make it possible to perform linkage mapping of fungal persistence directly in mammalian organs (Mullis et al. 2022).

In our previous work, we established such a system using a cross between the laboratory reference strain BY4716 (hereafter BY) and a haploid derivative of the clinical isolate 322134S (hereafter 3S) (Mullis et al. 2022). Using a pooled panel of 822 genotyped and barcoded haploid segregants, we measured relative persistence following tail-vein injection into C57BL/6J mice. Segregants showed reproducible, heritable differences in persistence across the brain, liver, kidneys, and spleen, and these patterns were broadly consistent across host sexes and between control mice and mice treated with dexamethasone to suppress immune function. Persistence in the brain was partly distinct from that in nonbrain organs. Linkage mapping identified 18 loci that fell into two classes. At general loci, the same parental allele favored persistence across organs. At antagonistically pleiotropic loci, an allele that favored persistence in the brain impaired persistence in nonbrain organs, or vice versa. High persistence in an organ was associated with both generally beneficial alleles and organ-appropriate alleles at antagonistically pleiotropic loci. Consistent with these trade-offs, many segregants persisted well in either the brain or nonbrain organs, but few persisted well in both. Thus, complex combinations of fungal polymorphisms both promoted and constrained persistence in different parts of the mammalian body.

Whether this genetic architecture generalized across host genotypes remained unknown because our previous study only included a single host genotype. This limitation matters because host genotype can substantially influence the outcome of fungal infection. For example, following intravenous infection with *Candida albicans*, inbred mouse strains differ markedly in survival, fungal burden, and tissue damage across organs (Ashman et al. 1996). A naturally occurring loss-of-function allele of complement component 5 (*C5*) is a major determinant of susceptibility in many inbred strains, but *C5*-independent loci and genetic-background effects also contribute (Tuite et al. 2005; Radovanovic et al. 2011). Such host genetic differences could change the relative performance of yeast genotypes, alter yeast allelic effect sizes, or reveal loci not detected in C57BL/6J mice. Alternatively, the genetic architecture of persistence could be dominated by yeast alleles whose effects remain similar across host backgrounds, or reflect a mixture of shared and host-dependent allelic effects. Testing these alternatives requires measuring the relative persistence of the same yeast genotypes across multiple host genetic backgrounds.

The eight founder strains of the Collaborative Cross and Diversity Outbred populations were selected to capture broad genetic diversity across the three major subspecies of *Mus musculus* and include both classical laboratory and wild-derived strains (Churchill et al. 2004; Churchill et al. 2012; Collaborative Cross Consortium 2012). We used seven of these founder strains to ask whether the genetic determinants of *S. cerevisiae* persistence vary across host genotypes. Following Mullis et al. (2022), we introduced a pool of barcoded BY × 3S segregants, along with both parental strains, into mice from each host genotype and then quantified relative persistence in the brain and liver two days later. We focused on these organs because our previous study showed that brain persistence was genetically distinct from persistence in the liver and other nonbrain organs. Persistence among the nonbrain organs was highly correlated, and yeast recovery was greatest on average from the liver. This design allowed us to quantify the relative contributions of yeast genotype and yeast genotype-by-host genotype interactions to relative persistence and compare linkage-mapping results among host genotypes.

We found that yeast genetic effects on relative persistence were largely shared across the tested host genotypes and that yeast genotype-by-host genotype interactions explained only a very small fraction of phenotypic variation in the analyzed samples.

## Methods

### Segregant panel

To identify regions of the *S. cerevisiae* genome associated with variation in persistence across organs and mouse strains, we used a pooled panel of haploid meiotic progeny (segregants) from the BY × 3S cross described in Mullis et al. (2022). The segregants were generated by crossing engineered haploid derivatives of the laboratory reference strain (BY) and the 322134S clinical isolate (3S), both carrying a genomic landing pad at the *YBR209W* locus (Liti et al. 2009; Levy et al. 2015; Liu et al. 2019; Matsui et al. 2022; Mullis et al. 2022). Specifically, *BY ho fcy1Δ flo8Δ flo11Δ ura3Δ YBR209W::pGal1-Cre-Lox71-Lox2272/71* and *3S ho fcy1Δ flo8Δ flo11Δ ura3Δ YBR209W::pGal1-Cre-Lox71-Lox2272/71* parental strains were used. Each segregant was represented by one or more unique, chromosomally integrated barcodes. The pool contained 872 barcoded segregants, as well as the BY and 3S parents (874 genotypes in total). Genomic genotype data were available for 816 of the segregants. To enable the estimation of broad-sense heritability, a subset of 160 segregant genotypes was replicated in the pool, with each replicate possessing a distinct barcode. Of these, 54 were represented by two distinct barcodes and 106 by three. BY and 3S were represented by seven and five distinct barcodes, respectively.

### Generation of the yeast pool

Segregants and parental strains were regrown individually and pooled before each experimental batch. Cryopreserved segregants and parental strains were individually inoculated into separate wells of 96-well deep-well plates containing 800 µL of YPD broth supplemented with 0.8 µL of a 200 mg/mL G418 stock per well. Following incubation at 30°C for 2 days, a yeast pool was generated by combining 500 µL from each culture. The pool was diluted to a concentration of 10⁸ cells/mL. A fraction of the pooled culture was pelleted to serve as the timepoint 0 (T0) control and stored at −20°C for later DNA extraction. The remaining suspension was aliquoted into 42 tubes, each containing the inoculum for a single mouse.

### Mouse experiments

All mouse procedures were approved by the University of Southern California’s Institutional Animal Care and Use Committee (protocol #21663). Mice were obtained from The Jackson Laboratory and were eight weeks old at injection. Mice were housed under a 12h light/dark cycle with food and water available *ad libitum*. Two days after injection, mice were euthanized by cervical dislocation.

The experiment was conducted in three batches using seven founder strains of the mouse Collaborative Cross and Diversity Outbred populations: 129S1/SvImJ, A/J, C57BL/6J, CAST/EiJ, NOD/ShiLtJ, NZO/HlLtJ, and PWK/PhJ (Churchill et al. 2004; Churchill et al. 2012; Collaborative Cross Consortium 2012). The eighth founder strain, WSB/EiJ, was excluded from the study because handling difficulties prevented successful tail-vein injection. For each batch, we injected three males and three females per strain (totaling 42 injections). Due to logistical constraints, only 14 individual mice could be injected on a given day. To balance host genotype and sex across injection days, we injected one male and one female from each of the seven strains on each day, with injection order randomized within each day. Mice were inoculated with 100 µL of the pool via tail-vein injection. Immediately following the injections, aliquots of the remaining pooled culture were plated on YPD + G418 to enable correction of differences among yeast strains in growth on selective medium. After 2 days at 30°C, yeast were scraped from the control plates, pelleted, and stored at −20°C for subsequent DNA extraction.

The brain and liver were collected from each mouse 2 days post-injection. Mice were euthanized via cervical dislocation and wiped down with 70% ethanol. We dissected the entire brain and liver from each animal, placing one brain hemisphere and two liver lobes into separate Qiagen PowerBead Tubes (Metal 2.38 mm). Organ processing and plating followed Mullis et al. (2022). Briefly, 1 mL of 1x TrypLE Select Enzyme (Thermo Fisher 12563029) was added to each tube, and samples were incubated at 37°C for 5 minutes before homogenization at 30 Hz for 3 minutes using a Qiagen TissueLyser II. The resulting homogenates were plated onto YPD agar supplemented with 200 µg/mL G418 and incubated at 30°C for 2 days to permit yeast colony formation. After 2 days of growth on the YPD + G418 agar, 10 mL of water was added to the surface of each plate and colonies were scraped and pelleted. The supernatant was then removed. These pellets were stored at −20°C for later DNA extraction. Samples with low yeast recovery based on visual inspection of colony growth on the plates were excluded from the study; this resulted in the removal of 23 samples from batch 1, 12 samples from batch 2, and 7 samples from batch 3.

### Barcode amplification

Genomic DNA was extracted from each sample pellet using the Zymo Quick-DNA Fungal/Bacterial Miniprep Kit and quantified via a Qubit high-sensitivity assay. Barcode sequencing libraries were then generated using a two-step PCR strategy. We amplified the barcoded genomic region using Phusion High-Fidelity DNA Polymerase (New England Biolabs) and custom primers containing both Unique Molecular Identifiers (UMIs) and sample-specific multiplexing tags (4 cycles). Multiplexing tags were designed to have a Hamming distance of 4 from one another. The primers were as follows, where N denotes a random nucleotide contributing to the UMI and X denotes a position in a known sample-specific multiplexing tag:

Forward:

5’-AATGATACGGCGACCACCGAGATCTACACNNXXXXNNACACTCTTTCCCTACACGACGCTCTTCCGATCTACGAAGTTATT GCGCGGTGA-3’

Reverse:

5’-CAAGCAGAAGACGGCATACGAGATNNXXXXNNGTGACTGG AGTTCAGACGTGTGCTCTTCCGATCT-3’

The 4-cycle PCR reaction was performed using the following steps:

1) 98°C, 3:00 min
2) 98°C, 30 s
3) 54°C, 30 s
4) 72°C, 2:00 min
5) Repeat steps 2 through 4 for a total of 4 cycles
6) 72°C, 5:00 min
7) 4°C, hold indefinitely

Afterwards, samples were treated with Exonuclease I (NEB) to degrade primers. Following purification with the Qiagen MinElute 96 UF PCR Purification Kit, a second PCR (24 cycles) was performed with the universal Illumina TruSeq Forward and Reverse primers to further amplify the barcode libraries. The 24-cycle PCR was performed using the following steps:

1) 98°C, 3:00 min
2) 98°C, 30 s
3) 54°C, 30 s
4) 72°C, 2:00 min
5) Repeat steps 2 through 4 for a total of 24 cycles
6) 72°C, 5:00 min
7) 4°C, hold indefinitely

Prior to sequencing, the resulting libraries were pooled and size-selected (∼220 bp) via electrophoresis and gel extraction.

### Quantifying barcode frequencies

Barcode amplicon libraries from each DO experiment were sequenced on separate lanes of an Illumina NovaSeq X Plus 10B flow cell (2 × 150 bp paired-end, 30% PhiX spike-in; Novogene), generating 970 Gb (3,233 M read pairs) in total. Paired-end reads were first filtered for quality; read pairs were retained only if the average Phred quality score of the first 35 base pairs in both the forward and reverse reads exceeded 30. The reads for different samples were then demultiplexed using a custom Python (version 3.11.9) script. The unique molecular identifiers (UMIs) and genotype barcodes (GenoBCs) were then extracted from the reads (Kivioja et al. 2012). To correct for errors introduced by PCR and sequencing, all unique GenoBC sequences observed across the dataset were aggregated and clustered with Bartender v1.1 using its default maximum merging distance of 2 and seed length of 5, with the statistical merging test disabled (-z −1) so that clusters were merged on sequence distance alone (Zhao et al. 2018). We identified a consensus sequence for each barcode cluster and matched it to known barcodes to identify the corresponding yeast genotype. Barcodes not matching any reference entry were discarded. Within each sample, reads sharing identical corrected GenoBCs and UMIs were collapsed into a single count. The final output reported the deduplicated count of each barcode in each sample.

### Quantifying persistence

Raw barcode counts were normalized within each sample by dividing each individual barcode count by the mean barcode count, yielding normalized barcode counts. Raw yeast persistence was then calculated as the relative change in normalized barcode count between the baseline (T0) and experimental endpoint (T1) using the formula: (T1 – T0)/T0. To correct for differences among yeast strains in on-plate growth, persistence measurements from growth controls were used as a reference. For each organ sample, a linear regression was fitted with raw barcode persistence values as the response and the corresponding growth-control values as the predictor. The residuals from this regression were retained as bias-corrected persistence phenotypes, replacing the raw persistence values for all downstream analyses (Mullis et al. 2022). Barcodes that were not detected in a sample, or in the corresponding T0 sample, were treated as missing for that sample rather than assigned zero persistence.

### Filtering samples by heritability

The subset of segregants represented by multiple barcodes was used to test for a significant genetic contribution to yeast persistence and estimate the heritability of yeast persistence in each organ sample. For each sample, a one-way ANOVA was performed testing persistence as a function of yeast genotype. We applied a Bonferroni-corrected significance threshold of P < 0.05/219 across the 210 organ samples and 9 homogenization controls, retaining only those samples that met this threshold. Broad-sense heritability (H^2^) was estimated for every organ sample by fitting a linear mixed model persistence ∼ genotype + error using the *lme4* package (version 1.1-37) in R (Bates et al. 2015). H^2^ was calculated as the proportion of total variance attributable to the genotype random effect. Samples were retained and used in subsequent analysis only if they met the significance threshold and had H^2^ > 0.2. Seven further samples were removed after inspection of sample-level clustering. Six were essentially uncorrelated with the retained samples of the same organ (mean r = −0.02 to 0.08, compared with at least 0.15 for every retained sample and a mean of 0.50; Supplementary Figure 1), and five of these also had variances far outside the range of the retained samples. The seventh, a PWK/PhJ liver sample (mean r = 0.21), was removed because it clustered with one of these outliers. Because its correlation with retained samples overlapped the lowest retained values, we confirmed that including it had a negligible effect on mean liver persistence (r = 0.99 across genotypes). While only two CAST/EiJ brain samples remained after heritability filtering, they were nonetheless included in the subsequent analysis; all other host-organ combinations retained a minimum of three samples following heritability filtering. For yeast genotypes represented by multiple barcodes, the first barcode replicate was selected for subsequent analyses, producing a single persistence value per genotype in each organ sample.

### Variance decomposition and across host analyses

To evaluate the consistency of yeast genotype effects across mouse backgrounds, linear models were fit separately for each organ in R (4.4.0):

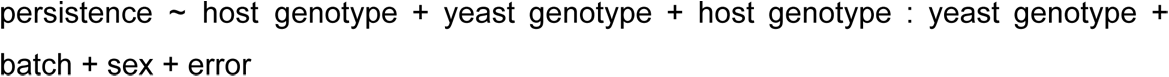

Batch, sex, host genotype, and yeast genotype were categorical variables corresponding to the experimental batch (1, 2, or 3), the sex of the injected mouse (M or F), the host genotype (129S1/SvImJ, A/J, C57BL/6J, CAST/EiJ, NOD/ShiLtJ, NZO/HlLtJ, or PWK/PhJ), and the individual yeast genotype, respectively. Batch and sex were included to account for technical and biological variation.

The model was fit in two ways. First, all terms were treated as fixed effects (lm) to test the main effects and the host genotype-by-yeast genotype interaction, using sequential (Type I) sums of squares in the order listed. Second, the same terms were treated as random effects (lmer, *lme4*; REML) to partition the total phenotypic variance. Variance components were expressed as percentages of the total phenotypic variance.

Not every yeast genotype was measured in every sample, and some yeast genotype–host genotype combinations were not observed. Both models were fit to the observed measurements only, without imputation. For each observed yeast genotype–host genotype combination, the fixed-effects model gives a fitted value: the model’s predicted persistence of that yeast genotype in that host genotype at the reference batch and sex. Because the model includes the full interaction term, this equals the mean of that yeast genotype’s measurements in that host genotype after adjusting for batch and sex. Combinations that were not observed have no fitted value and were excluded from the interaction tests and from the correlations between host genotypes, which used the yeast genotypes observed in both hosts of each pair.

From the fully fixed-effects model, we estimated an interaction effect for each yeast genotype in each host genotype under sum-to-zero coding: the fitted value of that yeast genotype in that host genotype minus the value predicted by the host genotype and yeast genotype main effects. The standard error of each interaction contrast was calculated from the fitted model’s coefficient variance–covariance matrix. Each effect was then tested against zero with a two-sided *t*-test on the residual degrees of freedom of the model (21,799 in brain and 53,183 in liver), and P values were Bonferroni-corrected across all yeast genotype–host genotype combinations within each organ. Fitted values were extracted from this model for every observed yeast genotype-by-host strain combination. These fitted values were arranged into a yeast genotype-by-host strain matrix. Pairwise Pearson correlations between host strains were computed to quantify the consistency of yeast genotype persistence values across mouse backgrounds.

### Parental comparisons

For each of the 14 host genotype–organ combinations, all genotypes were ranked by mean persistence, and parental ranks were expressed as percentiles. Parental mean persistence was compared across the 14 combinations with a one-sided paired *t*-test of whether 3S persistence exceeded BY persistence, with a one-sided Wilcoxon signed-rank test as a nonparametric check. Effect size was reported as Cohen’s *d* for the 14 paired host genotype–organ comparisons (the mean of the 3S − BY differences divided by their standard deviation).

### Procedure for quantitative trait locus mapping

Only the 816 segregant genotypes present in both the phenotype dataset and the genomic reference panel were used in linkage mapping. Linkage mapping was conducted for three phenotypes: brain persistence, liver persistence, and the brain-minus-liver difference. The brain-minus-liver difference phenotype aided in the detection of loci whose effects differ between organs.

Linkage mapping was performed with an iterative forward search. This procedure was used for all scans within each host genotype and across all host genotypes described below. The persistence phenotype for each yeast genotype was defined as the mean persistence across all samples that passed quality checks for a given organ, either within a host genotype or across all hosts, as specified below. The brain-minus-liver phenotype was the mean brain persistence minus the mean liver persistence of each yeast genotype. Each genome-wide scan was conducted in each mapping round using one-way ANOVA at every SNP marker in R with the lm() and summary.aov() functions, and variance explained was the R^2^ of this model.

Within each mapping round, the most significant SNP on each chromosome was declared a locus if it exceeded a permutation-derived significance threshold and was more than 100 kb from any previously identified peak on that chromosome. The significance threshold was calculated separately for every scan (each phenotype, host genotype or aggregate mapping, and mapping round) from 1,000 permutations. In each permutation, the phenotype values were randomly shuffled among segregants while each segregant’s genotypes were kept intact, and the smallest P value across all markers was recorded. The 5th percentile of these 1,000 minimum P values was used as the genome-wide significance threshold (α = 0.05) (Churchill and Doerge 1994). In subsequent mapping rounds, the phenotype was adjusted for all previously detected loci by regressing it on the genotypes at their peak markers. The residuals from this regression were tested for association at every marker in place of the original phenotype. In the permutations for these rounds, each shuffled phenotype was adjusted in the same way before scanning. This process continued until no new loci were detected, allowing multiple loci to be identified on the same chromosome across successive rounds. Peak positions, variance explained, and support intervals, defined by a 2-unit drop in −log₁₀(P) from the peak, were recorded.

We first performed linkage mapping independently for each host genotype (129S1/SvImJ, A/J, C57BL/6J, CAST/EiJ, NOD/ShiLtJ, NZO/HlLtJ, and PWK/PhJ). For this host-specific approach, only the mouse samples belonging to that strain were used to compute per-yeast-genotype mean persistence values. Mapping continued until no additional locus exceeded the permutation threshold; no strain-specific scan required more than five rounds. The number of samples passing quality filters varied by host (129S1/SvImJ Brain: 4, Liver: 12; A/J Brain: 7, Liver: 10; C57BL/6J Brain: 4, Liver: 8; NZO/HlLtJ Brain: 3, Liver: 9; PWK/PhJ Brain: 5, Liver: 10; NOD/ShiLtJ Brain: 9, Liver: 14; CAST/EiJ Brain: 2, Liver: 10).

In addition to the strain-specific mapping, linkage mapping was performed using each yeast genotype’s mean persistence across all retained samples for a given organ (34 brain samples or 73 liver samples). Mapping was performed in the same iterative permutation-based mapping framework that was applied in the strain-specific mapping. As in the strain-specific mapping, scans continued until no additional locus exceeded the permutation threshold; no aggregate scan required more than three rounds.

### Defining and matching loci

Detections were combined into individual loci by the overlap of their support intervals. For the mapping across all host genotypes, detections from all mapping rounds and from the brain, liver and brain-minus-liver scans were merged if their support intervals overlapped by at least 1 kb. Each merged locus spans the union of its merged intervals, and its peak marker is that of its most significant detection (highest −log₁₀(P)); this marker was used for all allelic-effect, variance-explained and additive-model estimates. For the mapping within each host genotype, detections within a host genotype were first correspondingly merged if their intervals overlapped or lay within 1 kb of each other. The resulting host-level loci were then merged across host genotypes if their intervals overlapped by at least 1 kb, giving 24 within-host loci. A within-host locus was counted as detected in the aggregate mapping if its interval overlapped an aggregate locus by at least 1 kb. An aggregate locus was counted as previously detected if its interval overlapped a locus reported by Mullis et al. (2022) by at least 1 kb.

### Detection breadth and variance explained

A locus could be detected more than once within a host genotype, in different phenotype scans or mapping rounds. Variance explained was the R^2^ of the peak marker for the phenotype analyzed in the mapping round in which the locus was detected. For each locus detected within a host genotype, we therefore took the largest R^2^ among its detections in that host and averaged these values across the host genotypes in which the locus was detected. Loci were grouped by the number of host genotypes in which they were detected (1, 2, 3, 6, or 7). Mean variance explained was compared among groups by one-way ANOVA, followed by Tukey’s HSD test for pairwise comparisons between groups.

### Allelic effects and locus classes

For each aggregate locus, the estimated allelic effect in each organ was the difference in mean persistence between segregants carrying the 3S allele and those carrying the BY allele at the peak marker (3S − BY). A locus was classified as antagonistically pleiotropic if its estimated brain and liver effects had opposite signs, and as general if they had the same sign. Within each class, effect magnitudes (absolute values) in brain and liver were compared with paired *t*-tests across loci.

### Beneficial-allele counts

For each segregant, we counted the alleles associated with higher persistence in each organ at each locus class. At general loci, the allele associated with higher mean persistence in both organs was counted as beneficial. At antagonistically pleiotropic loci, the brain-beneficial and liver-beneficial alleles were counted separately. Organ-specific totals were the general count plus the count of alleles beneficial for that organ at antagonistically pleiotropic loci. Associations between allele counts and persistence were tested with Spearman rank correlation across the 816 genotyped segregants.

### Additive model of the 17 aggregate loci

We performed linkage mapping for persistence averaged across all mouse hosts in brain, liver and brain-minus-liver. From this scan we identified 17 aggregate loci. For each organ, fitted persistence values from the fixed-effects model were averaged across the host genotypes in which each segregant was observed. Brain and liver persistence were each regressed on the genotypes at the peak markers of the 17 aggregate loci across the 816 genotyped segregants. Parental persistence was predicted from the parents’ genotypes at these markers; the parents were not included in the fit. Ninety-five percent confidence intervals for model R^2^, the predicted parental values, and the predicted 3S − BY difference were calculated as bootstrap percentile intervals from 1,000 resamples of segregants with replacement, with the 17 loci held fixed. We also reported the proportion of bootstrap replicates in which the predicted 3S value exceeded the predicted BY value.

### Genetic variance explained by the aggregate loci

To estimate the proportion of genetic variance explained by the 17 aggregate loci, two models were fit to all individual persistence measurements for an organ from the 816 genotyped segregants. Broad-sense heritability (H^2^) was the yeast genotype variance divided by the total variance in the random-effects model persistence ∼ yeast genotype + error. The variance explained by the loci (R^2^) was taken from the linear model persistence ∼ genotypes at the peak markers of the 17 aggregate loci + error. Under this genotype-only model, the proportion of between-genotype variance explained was approximated by R^2^/H^2^ (Supplementary Table 12).

### Use of large language models

Large language models were used to assist with code development, statistical checks, and manuscript editing. Claude (Anthropic; Claude Code with Claude Opus models; February–October 2026) was used to help write and refactor the R and Python analysis code, to check reported statistics against the analysis outputs, and to edit manuscript text. ChatGPT (OpenAI; Codex with Sol and Astra models, February–October and September–October 2026, respectively) was used mainly to edit manuscript text and figures and was not used to write analysis code. All code was reviewed, all results were verified against the analysis outputs, and we take full responsibility for the content.

## Results

### Analysis of yeast persistence across founder strains of the mouse Diversity Outbred population

We attempted tail-vein injection of the pool of 874 barcoded yeast genotypes, which included BY, 3S, and 872 unique BY × 3S segregants, into mice from the eight founder strains of the mouse Collaborative Cross and Diversity Outbred populations: 129S1/SvImJ, A/J, C57BL/6J, CAST/EiJ, NOD/ShiLtJ, NZO/HlLtJ, PWK/PhJ, and WSB/EiJ. Because WSB/EiJ mice were difficult to handle, we were unable to inject that strain and proceeded with the remaining seven mouse strains. Across the 126 mice injected with the segregant pool, we obtained 252 organ samples (126 brain and 126 liver). After homogenizing each organ sample and selectively plating the homogenates to enrich for yeast, we retained 210 organ samples (100 brain and 110 liver) that had sufficient yeast recovery for further analysis. We extracted DNA from the recovered yeast and performed barcode sequencing.

To identify samples in which segregant genotype had a significant effect on persistence measurements, we performed a one-way ANOVA for each sample using the 160 segregants that were represented by two or three distinct barcodes. Using a Bonferroni-corrected family-wise threshold of α = 0.05, 39 brain and 77 liver samples showed significant genotype effects. Among these samples, broad-sense heritability (H^2^) ranged from approximately 0 to 1 in the brain and from 0.20 to 0.82 in the liver (Figure 2a). Two of these brain samples had H^2^ values ≤ 0.005 and were excluded from further analysis, leaving 37 brain and 77 liver samples. In addition, sample-level clustering identified 3 brain and 4 liver samples as outliers; these samples were also excluded (Supplementary Figure 1). In total, 34 brain and 73 liver samples were retained for all downstream analyses (Supplementary Table 1).

**Fig. 1.**
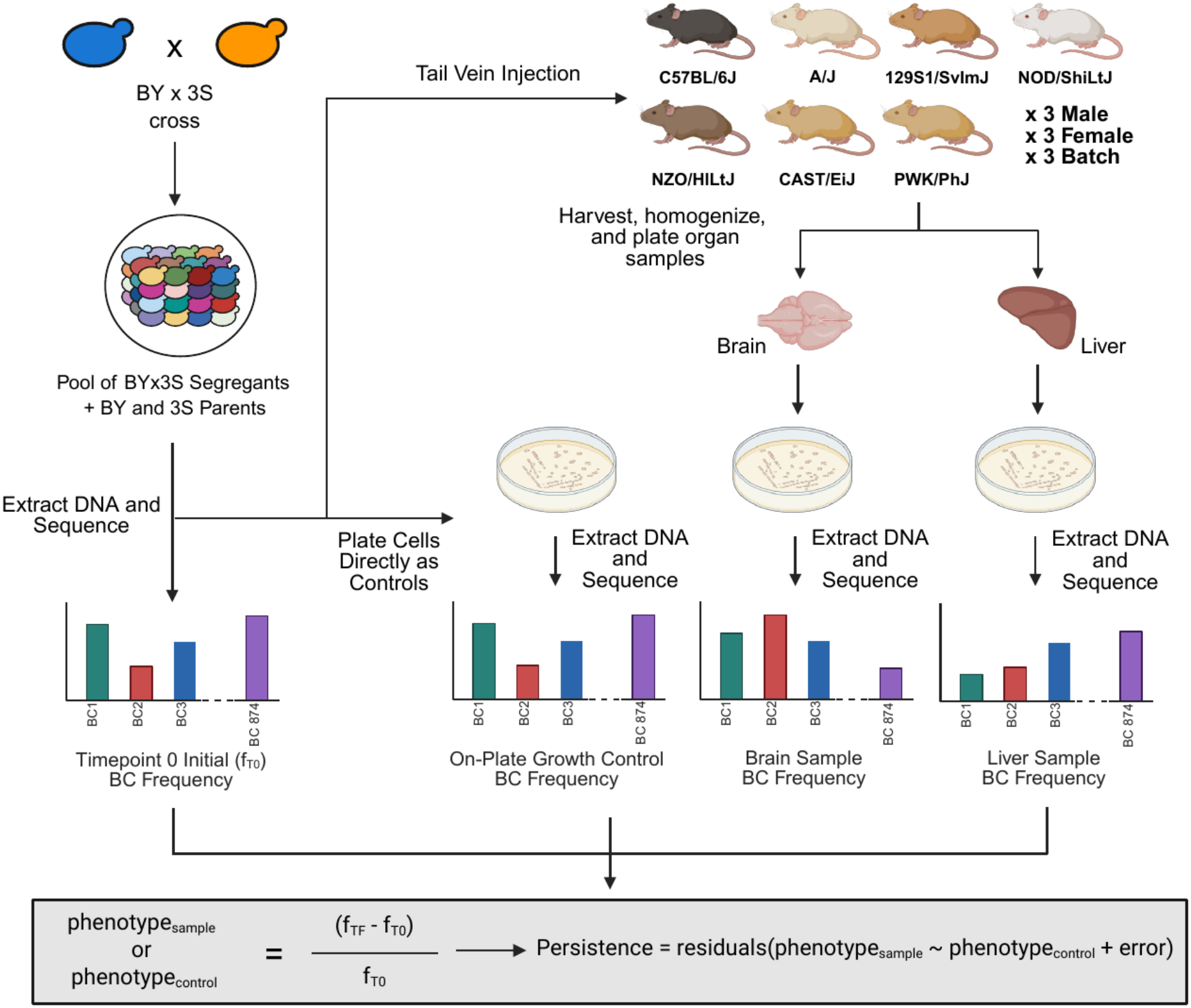
Experimental design. Barcoded BY×3S segregants and both parents were pooled and injected via the tail vein into mice from seven Collaborative Cross and Diversity Outbred founder strains (nine males and nine females per strain across three batches). Brain and liver were harvested two days after injection, and their homogenates were plated to recover yeast. Barcodes from the recovered yeast were sequenced alongside a time-zero pool and on-plate growth controls. Persistence was defined for each barcode as the residual from regressing the relative change in normalized barcode count in an organ sample on the corresponding change in its matched growth control. Alt text: Schematic of the experiment. A cross between two yeast strains, BY and 3S, produces barcoded offspring, which are pooled with both barcoded parents and injected into the tail vein of mice from seven inbred strains, three males and three females per strain in each of three batches. Brains and livers are harvested, homogenized and plated, and the barcodes of the recovered yeast are sequenced and compared with the injected pool and with growth controls.

**Fig. 2.**
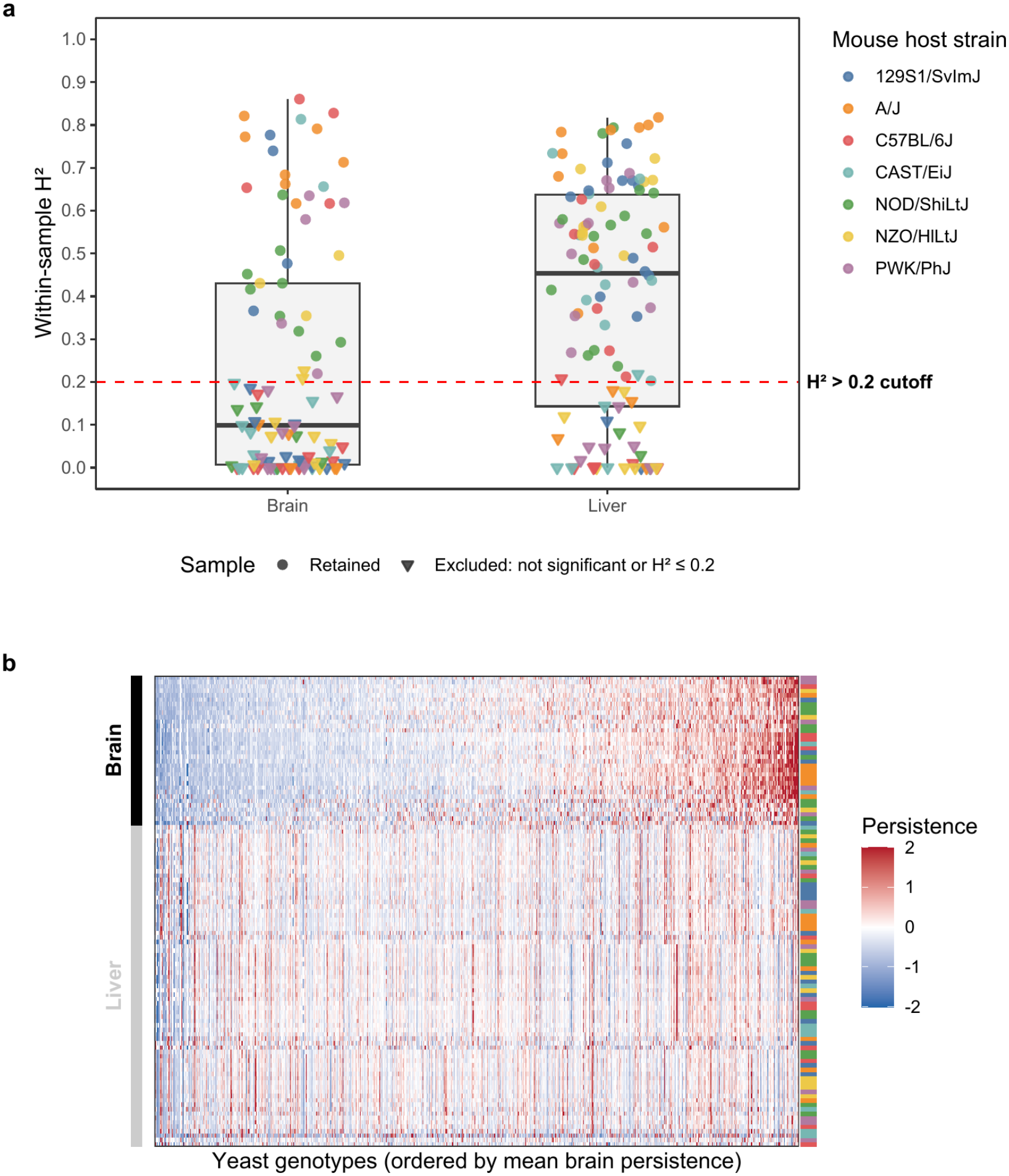
Heritability and structure of persistence across samples. **a)** Within-sample broad-sense heritability (H^2^) for the 203 brain and liver samples, excluding the seven samples removed as outliers, colored by mouse host genotype. H^2^ was estimated from the segregants represented by multiple barcodes. Boxes show the median and interquartile range; whiskers extend to the most extreme values within 1.5 times the interquartile range. The dashed red line marks H^2^ = 0.2; samples at or below it were excluded from downstream analyses. Circles, retained samples; triangles, samples excluded because their genotype effect was not significant after Bonferroni correction or their H^2^ was 0.2 or less. **b)** Persistence (color scale truncated at ±2) for 874 yeast genotypes across the 107 retained samples (34 brain and 73 liver). Genotypes (columns) are ordered by mean brain persistence; samples (rows) are clustered within organ (Euclidean distance, complete linkage). Left bar, organ; right bar, mouse host genotype, colored as in a). Alt text: Two panels. a) Box plots with overlaid points of within-sample heritability for brain and liver samples, colored by mouse strain; liver samples have higher heritability than brain samples, and a dashed red line at 0.2 marks the cutoff, with retained samples shown as circles and excluded samples as triangles. b) Heat map of persistence for 874 yeast genotypes (columns, ordered by mean brain persistence) across 107 samples (rows, clustered), with brain and liver samples separated; brain samples show a strong gradient from low to high persistence that liver samples largely lack.

### Mouse host genotype-by-yeast genotype interactions explain little phenotypic variance

The main goal of our study was to test whether the genetic architecture of yeast persistence in mice varies across host genotypes. Because the persistence measures generated by our pooled assay are relative, our data do not address differences in absolute fungal burden among host genotypes. Instead, we tested whether individual yeast genotypes performed better or worse in particular host genotypes by examining host genotype-by-yeast genotype interactions.

To identify such statistical interactions, we fit separate fixed-effects linear models for the brain and liver that included batch, sex, host genotype, yeast genotype, and the host genotype-by-yeast genotype interaction. In the models, batch, sex, and host genotype did not have significant main effects in either organ (P > 0.05). By contrast, yeast genotype was significant in both organs (P < 1 × 10⁻³⁰⁰), as was the host genotype-by-yeast genotype interaction (P ≤ 6.01 × 10⁻⁵⁹; Supplementary Tables 2 and 3).

We determined the variance explained by the batch, sex, host genotype, yeast genotype, and the host genotype-by-yeast genotype interaction terms using random-effects linear models. Yeast genotype explained 44.3% of the variance in brain persistence and 46.0% in liver persistence, whereas yeast genotype-by-host genotype interactions explained only 4.4% and 1.3%, respectively (Supplementary Tables 4 and 5). Host genotype, batch, and sex each explained 0.02% or less of the variance in either organ (brain: 0%, 0.002%, and 0%; liver: <0.001%, 0.02%, and <0.001%, respectively), and residual variance accounted for the remaining 51.3% in brain and 52.7% in liver. Consistent with the small contribution of host genotype-by-yeast genotype interactions, fitted persistence values for individual yeast genotypes were positively correlated between every pair of host genotypes (brain: mean r = 0.70, range 0.58–0.83; liver: mean r = 0.82, range 0.65–0.94; Supplementary Figure 2).

To identify significant host genotype-by-yeast genotype interactions, we estimated the interaction effect of each yeast genotype in each host genotype from the fixed-effects linear models for brain and liver. We then tested whether each effect differed from zero using two-sided *t*-tests of the model-derived interaction contrasts, applying a Bonferroni-corrected threshold of α = 0.05 across all combinations tested within each organ. We found 31 and 33 significant interactions in the brain and liver, respectively, among the 6,088 and 6,113 yeast genotype–host genotype combinations tested (Supplementary Tables 6 and 7). In the brain, these 31 interactions involved 22 segregants: two segregants were involved in three interactions, five in two, and 15 in one interaction each. In the liver, the 33 interactions involved 20 segregants: two were involved in five interactions, one in four, two in two, and 15 in one interaction each. Four segregants showed significant interactions in both organs. Every host genotype had at least one significant interaction with a segregant in one or both organs; C57BL/6J showed significant interactions only in the liver (Figures 3 and 4). In summary, host genotype-by-yeast genotype interactions explained only a small fraction of phenotypic variance, and significant interactions were detected in a small subset of segregants.

**Fig. 3.**
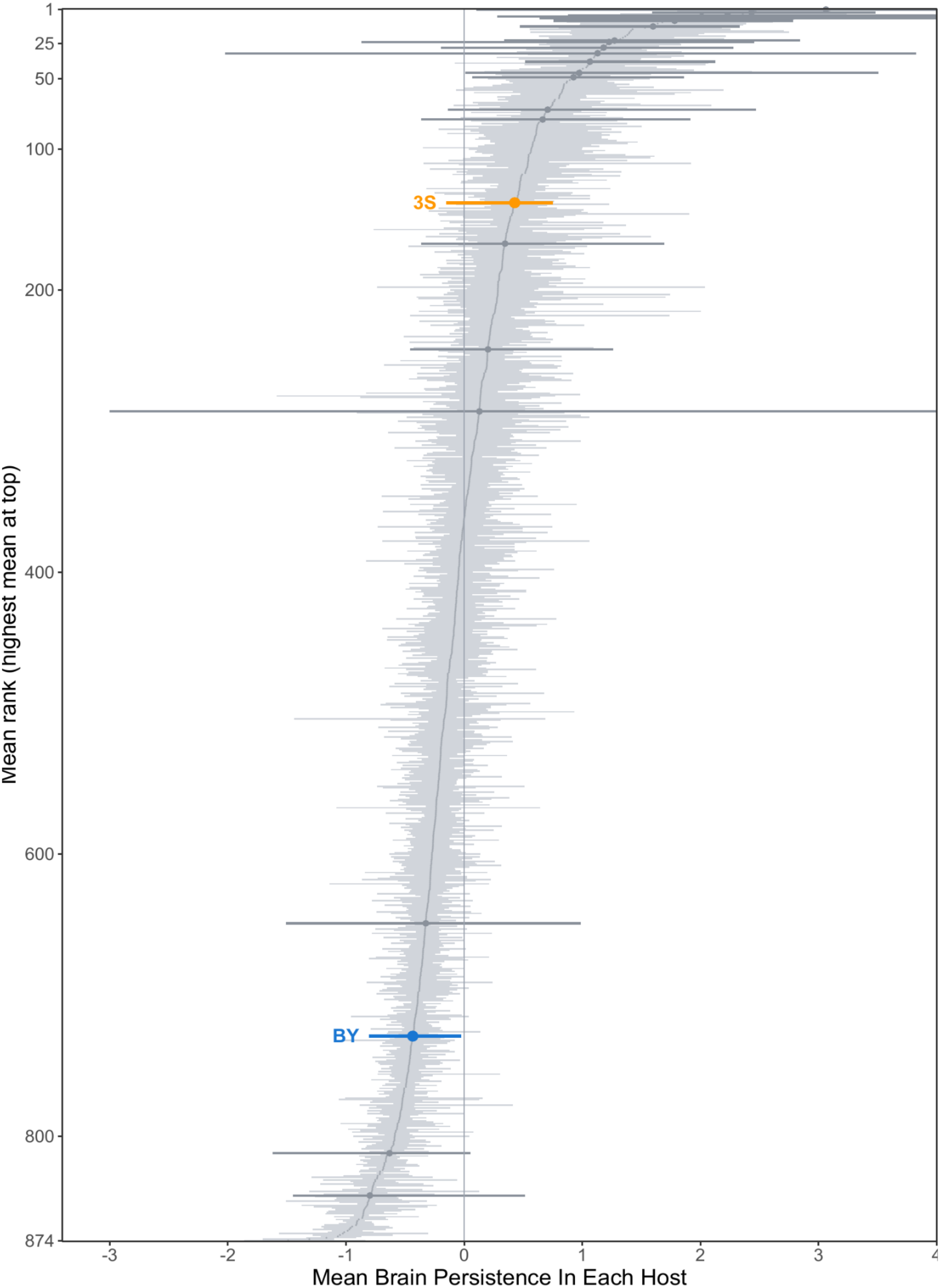
Brain persistence of each yeast genotype across host genotypes. Genotypes (n = 874, including both parents) are ranked by mean brain persistence, highest at top. Each light grey line spans the range of one genotype’s mean persistence values in the individual host genotypes, and grey dots mark its mean across host genotypes; lines show variation across host genotypes. BY (blue) and 3S (orange) are highlighted. Darker grey lines mark the 22 segregants with at least one significant host genotype-by-yeast genotype interaction in the brain after Bonferroni correction (adjusted P < 0.05; Supplementary Table 6). The x-axis is truncated for readability; segments extending beyond the limits are cut at the axis edge. Alt text: Horizontal line plot of 874 yeast genotypes ranked from highest (top) to lowest (bottom) mean brain persistence. Each light grey line spans one genotype’s mean persistence across the seven mouse strains, with a grey dot at its overall mean, forming a diagonal band. Darker grey lines mark the 22 segregants with a significant host-by-yeast interaction, most of which have unusually wide ranges. The parent 3S (orange) ranks near the top and BY (blue) near the bottom.

**Fig. 4.**
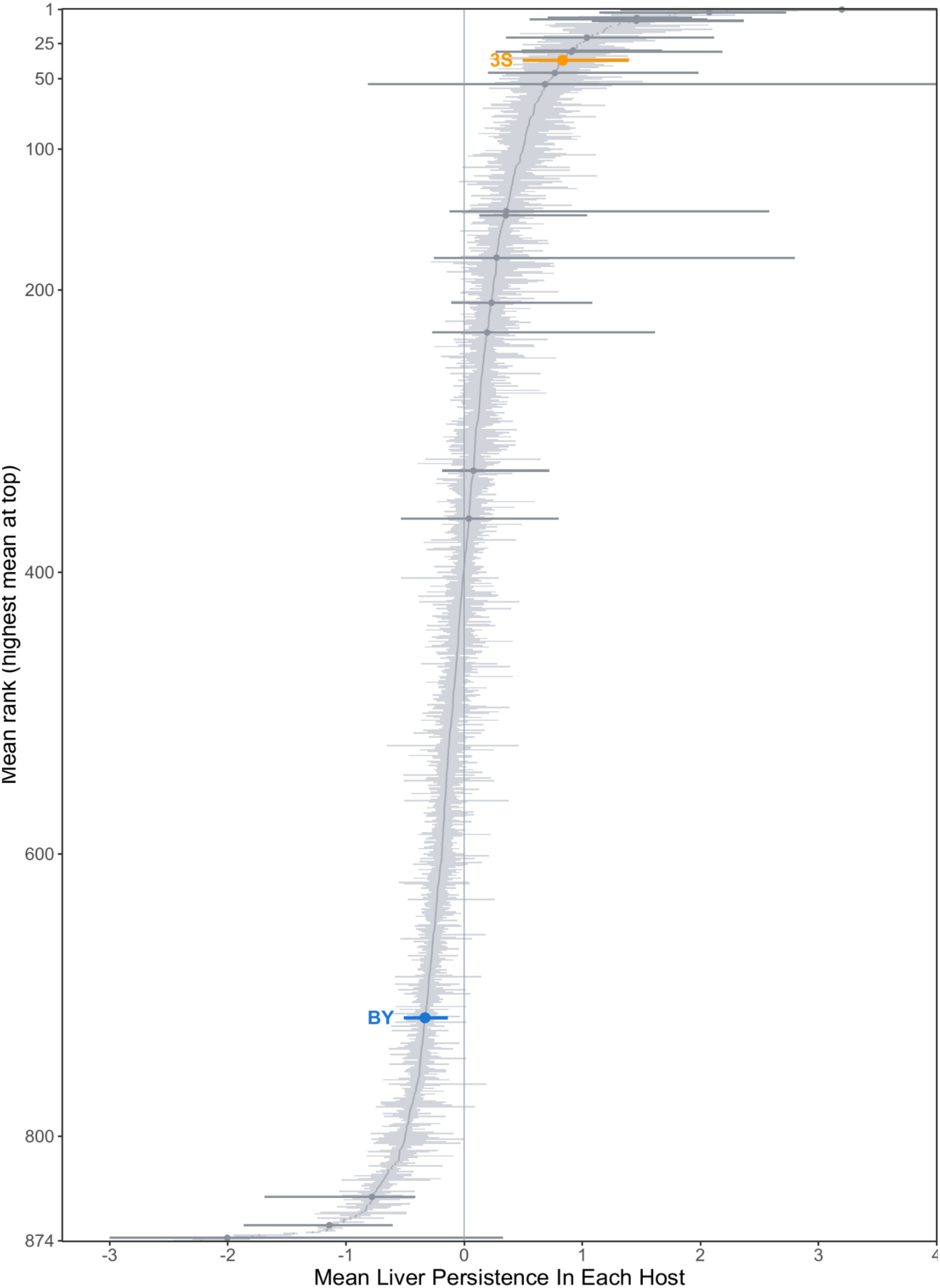
Liver persistence of each yeast genotype across host genotypes. Plotted as in Fig. 3, with genotypes ranked by mean liver persistence. Darker grey lines mark the 20 segregants with at least one significant host genotype-by-yeast genotype interaction in the liver after Bonferroni correction (adjusted P < 0.05; Supplementary Table 7). Alt text: The same plot as Figure 3 for liver persistence. The band of genotypes is narrower than in brain; darker grey lines mark the 20 segregants with a significant host-by-yeast interaction, and 3S (orange) ranks near the top while BY (blue) ranks low.

### Weak correlation between brain and liver persistence among segregants

Persistence measurements from individual organ samples are noisy; thus, we averaged persistence measurements for each segregant within each host genotype–organ combination (Figures 3 and 4). Mean fitted persistence, averaged across host genotypes, showed transgressive segregation in both organs, with segregant phenotypes extending beyond the parental values (Figure 5). Mean fitted brain and liver persistence were only weakly correlated across segregants (Pearson r = 0.15, P = 8.72 × 10⁻⁶; Figure 5), and most segregants that showed high persistence did so in only one of the two organs. Of the 164 unique segregants in the top 10% for persistence in at least one organ, 12 (7%) were in the top 10% for both.

**Fig. 5.**
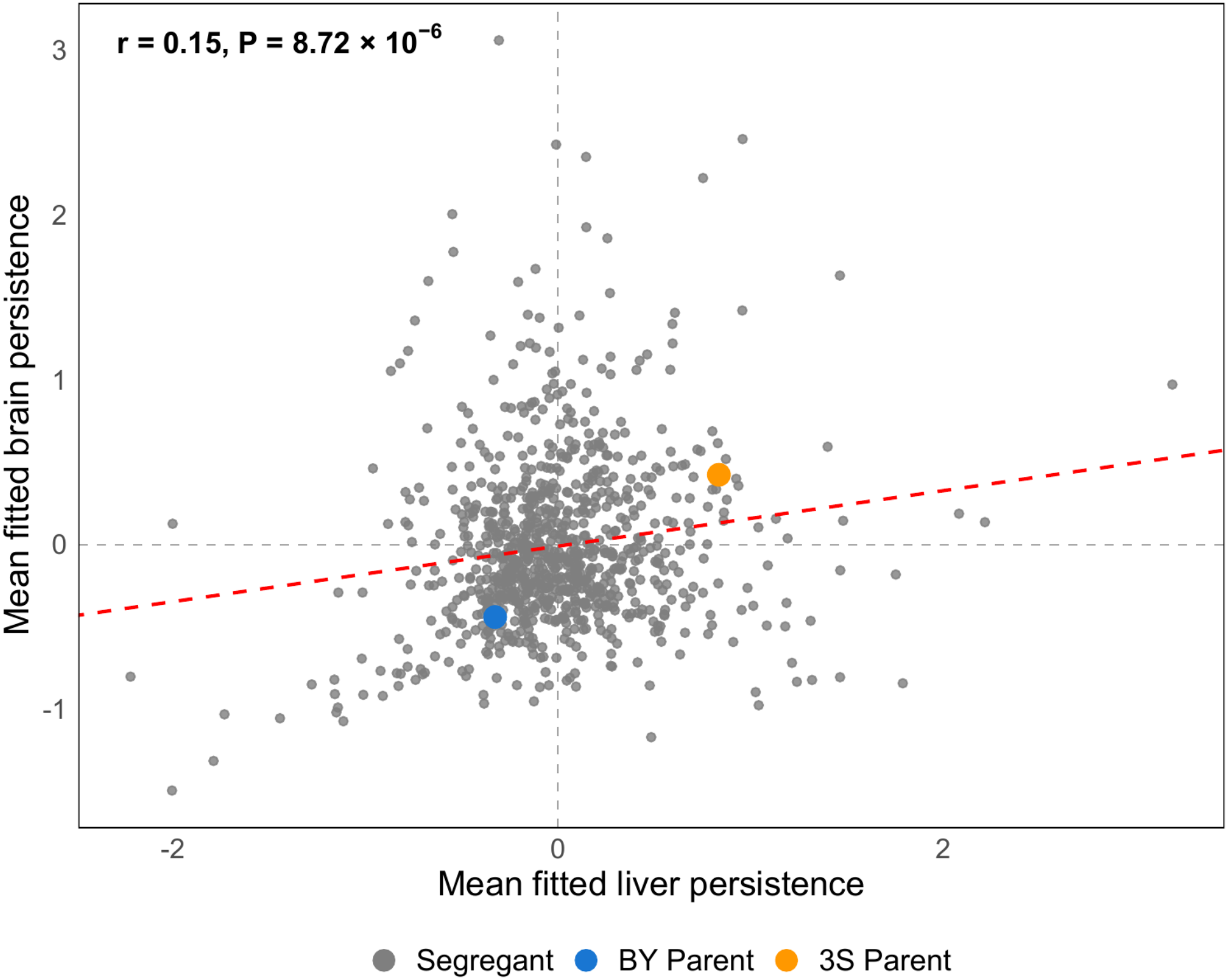
Mean fitted brain persistence (y-axis) against mean fitted liver persistence (x-axis) for each yeast genotype. Fitted values are from the fixed-effects model, averaged across the host genotypes in which the genotype was observed. Grey, segregants; blue, BY parent; orange, 3S parent. Dashed red line, least-squares fit across segregants (Pearson r = 0.15, P = 8.72 × 10⁻⁶; parents excluded). Dashed grey lines mark zero persistence in each organ. Alt text: Scatter plot of mean fitted brain persistence against mean fitted liver persistence for each yeast genotype. The grey segregant points form a diffuse cloud with a weak positive trend (dashed red line, Pearson r = 0.15). The 3S parent (orange) lies in the upper right and the BY parent (blue) in the lower left.

### 3S outperforms BY in the brain and liver across all hosts

Our previous study did not include both parental strains in the pooled assay. Here, the pool included barcoded BY and 3S strains, enabling direct comparison of parental persistence across host genotypes and organs. Across all host genotype–organ combinations, 3S exhibited higher mean persistence than BY (Figure 5; Supplementary Table 8). BY ranked in the lower half of genotypes in 13 of 14 comparisons (the exception was NZO/HlLtJ brain), whereas 3S ranked in the top 28% of genotypes in every comparison except the CAST/EiJ brain comparison (top 49%), which was based on only two samples. This separation was more pronounced in the liver, where 3S frequently ranked in the top decile and BY near the bottom quartile. A one-sided paired *t*-test across the 14 host–organ comparisons supported the higher mean persistence of 3S relative to BY (*t* = 12.01, df = 13, P = 1.03 × 10⁻⁸, Cohen’s d = 3.21). Consistent with this pattern, when persistence was averaged across host genotypes, 3S ranked in the top 15% (brain) and top 4% (liver) of genotypes, while BY was in the bottom 16% and 19%, respectively. Mean fitted persistence values from the fixed-effects model, averaged across host genotypes, were −0.44 in the brain and −0.33 in the liver for BY, compared with 0.42 and 0.84, respectively, for 3S (Figure 5). These results demonstrate a consistent persistence advantage of 3S over BY in both organs across host genotypes.

### Loci detected in only one host explain little variance

We performed linkage mapping for brain persistence, liver persistence, and the brain-minus-liver difference within each mouse host genotype using the 816 genotyped segregants (Supplementary Table 9). Within each host genotype, we combined results from these three linkage-mapping scans to identify distinct loci, because some loci were detected in more than one scan. Between 9 (PWK/PhJ) and 17 (A/J) unique loci were detected per host. Across host genotypes, these loci corresponded to 24 distinct loci. Seven were detected in all seven host genotypes, three in six, four in three, and two in two. The remaining eight were each detected in a single host genotype: four in CAST/EiJ and one each in A/J, C57BL/6J, NZO/HlLtJ, and NOD/ShiLtJ (Figure 6).

**Fig. 6.**
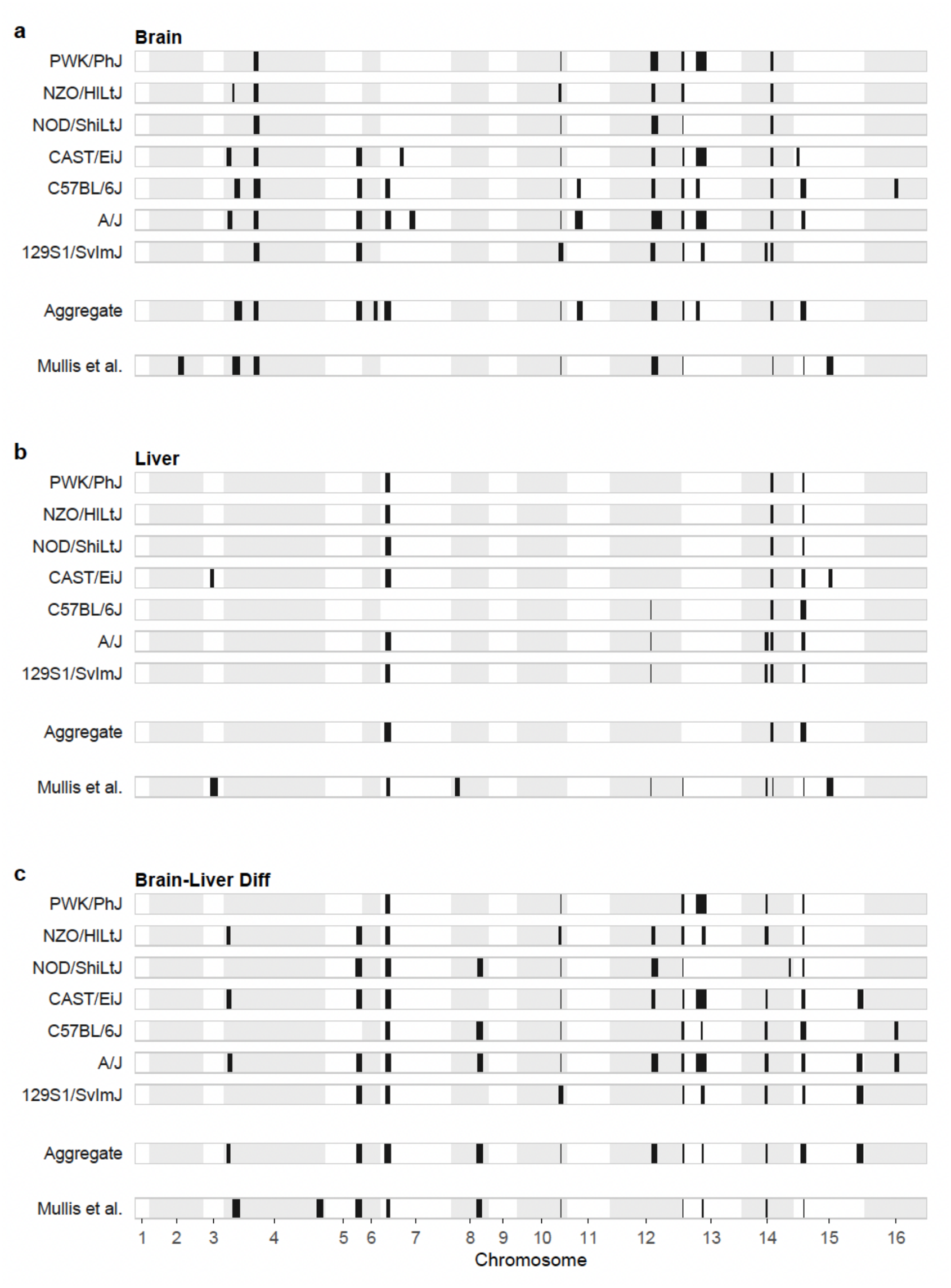
Genomic distribution of persistence loci by phenotype. **a)** Brain. **b)** Liver. **c)** Brain-minus-liver difference. Within each panel, the upper seven rows show loci mapped separately within each host genotype; below, separated by additional space, are loci from the aggregate mapping across all hosts and from Mullis et al. (2022). Black bars mark support intervals (2-unit drop in −log₁₀(P) from the peak; 816 genotyped segregants in the current study); alternating shading distinguishes the 16 chromosomes. For Mullis et al. (2022), panels b) and c) show loci mapped for combined nonbrain persistence and the brain-minus-nonbrain difference, respectively. Alt text: Three genome maps, for brain, liver and the brain-minus-liver difference, each spanning the 16 yeast chromosomes on the horizontal axis. In each map, the top seven rows show locus support intervals as black bars for each mouse strain, followed by one row for loci from all hosts combined and one row for loci from Mullis et al. (2022). Many intervals line up vertically across mouse strains, showing loci shared among hosts.

Loci detected in all seven hosts individually explained an average of 5.23% of the variance in the corresponding mapping phenotype within a host, with a maximum of 12.45% for a locus on chromosome X. Each of the eight single-host loci explained only 0.98–2.91% of the corresponding mapping-phenotype variance (mean 1.51%). Loci detected in more host genotypes tended to explain more variance, and mean variance explained differed significantly among detection-breadth groups (one-way ANOVA: F(4,19) = 6.67, P = 0.0016; Figure 7a). Loci detected in all seven host genotypes explained significantly more variance per host than loci detected in only one host genotype (Tukey HSD: Δ = 3.72 percentage points of variance explained, adjusted P = 8.1 × 10⁻⁴).

**Fig. 7.**
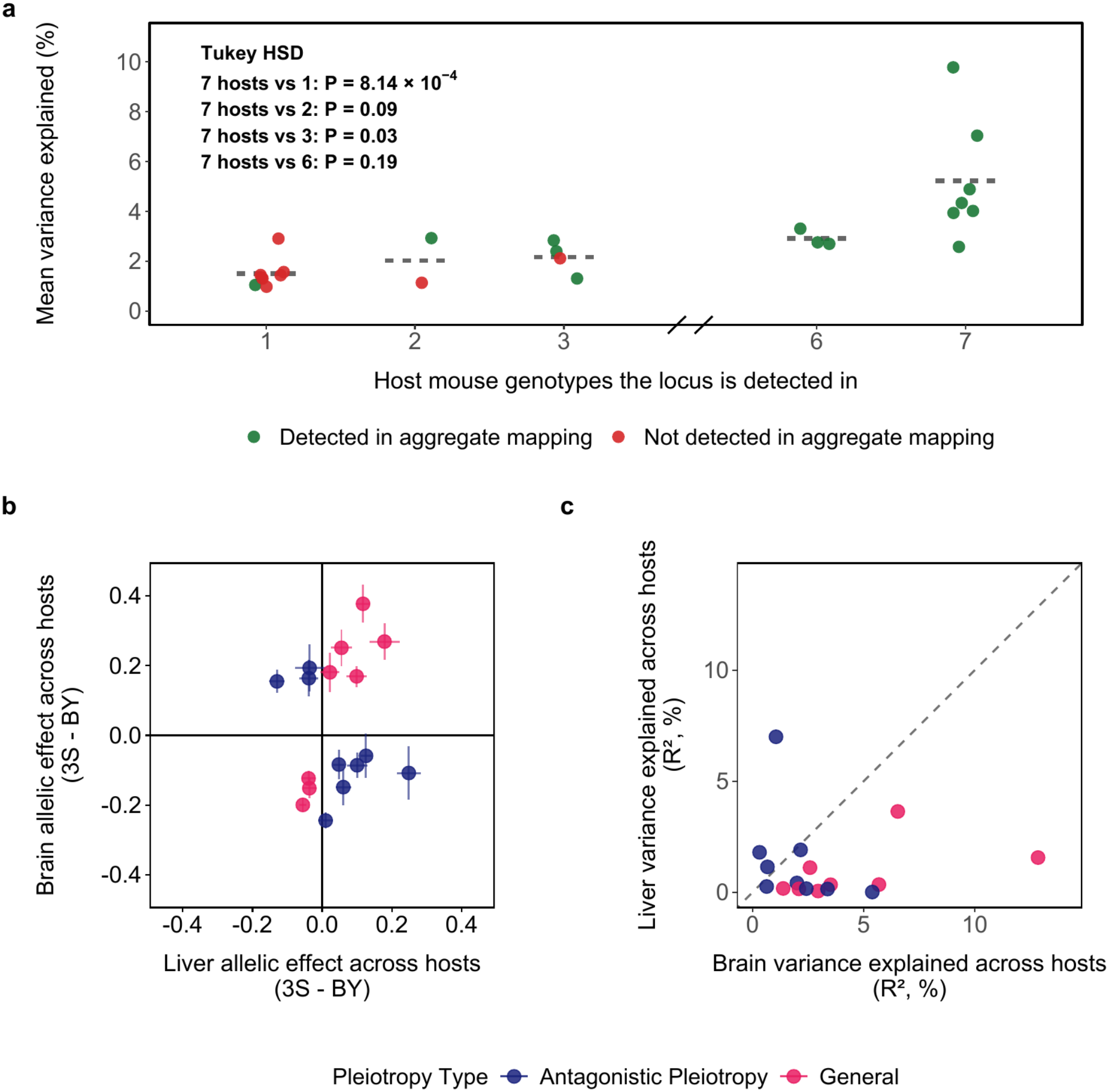
Detection breadth, pleiotropy and organ specificity of persistence loci. **a)** Mean variance explained per host for each of the 24 within-host loci (for each locus, the largest R^2^ among its detections in a host, averaged across the host genotypes in which it was detected), against the number of host genotypes in which it was detected. Green, loci also detected in the aggregate mapping; red, the 9 loci detected in within-host scans but not in the aggregate mapping. Dashed crossbars, means of each detection-breadth group. The x-axis is broken between 3 and 6 because no locus was detected in exactly four or five hosts. Black text, Tukey HSD post hoc P values for the 7-host group against each other group. **b)** Brain (y-axis) against liver (x-axis) allelic effect (3S − BY) of the 17 aggregate loci, estimated from mean persistence across all retained samples; crosshairs show the standard deviation of the seven host-specific allelic effects, each estimated from mean persistence within one host genotype. **c)** Liver (y-axis) against brain (x-axis) variance explained (R^2^) for the same loci; dashed line, y = x. Colors in b) and c) denote locus class: pink, general; navy, antagonistically pleiotropic. Alt text: Three panels. a) Dot plot of mean variance explained by each of 24 loci against the number of mouse strains in which it was detected (1, 2, 3, 6 or 7); loci found in all seven strains explain the most variance, and loci found in one strain explain little. Green points are loci also found in the combined mapping and red points are not. b) Scatter plot of brain against liver allelic effects for 17 loci, colored by class: general loci (pink) lie in the same-sign quadrants and antagonistically pleiotropic loci (navy) in the opposite-sign quadrants, with crosshairs showing variation across hosts. c) Scatter plot of liver against brain variance explained for the same loci with a dashed one-to-one line; 14 of the 17 loci explain more variance in brain than in liver.

We also compared host-level patterns of single-host locus detection and significant host genotype-by-yeast genotype interactions. However, there was no consistent correspondence between these patterns (Supplementary Table 10).

### A common set of loci explains variation in yeast persistence across host genotypes

Because many of the loci were detected across different hosts, we sought to increase our statistical power in linkage mapping by averaging each segregant’s persistence across all retained samples within each organ and mapping these means and their difference. Linkage mapping of these aggregate phenotypes identified 17 distinct loci (Figure 6; Supplementary Table 11). These loci collectively explained 45.32% and 19.07% (adjusted R^2^) of the variance in mean brain and liver persistence, respectively.

Using individual persistence measurements rather than means, we separately estimated the proportion of between-genotype variance explained by the 17 loci. Dividing the R^2^ values from this analysis by the corresponding genotype-only heritability estimates (H^2^ = 0.45 in brain and 0.46 in liver) gave approximate estimates of 49% and 22%, respectively (Supplementary Table 12).

The three loci explaining the most phenotypic variance mapped to chromosomes X, XIV, and XV. The chromosome X locus explained the most brain-persistence variance (R^2^ = 12.9%), the chromosome XV locus explained the most liver-persistence variance (R^2^ = 7.0%), and the chromosome XIV locus contributed substantially in both organs (brain R^2^ = 6.5%, liver R^2^ = 3.6%; Figure 7c). Of the 17 loci detected in the current study, 12 were also identified in our prior work, which found 18 distinct loci (Mullis et al. 2022; Figure 6). The support intervals for the 17 loci ranged from 10.8 to 95.3 kb, limiting resolution of the causal genes and mechanisms (Supplementary Table 11).

### General loci have larger effects in the brain than the liver

In our previous work, we found that some loci had effects in the same direction in the brain and liver (“general”), whereas others had opposite effects in the two organs (“antagonistic pleiotropy”). The same pattern was evident in the aggregate data. Eight of the 17 loci in the current study were classified as general, and 9 were classified as antagonistically pleiotropic, based on the directions of their estimated effects. The BY and 3S alleles favored persistence at three and five of the general loci, respectively (Figure 7b).

General loci had significantly larger absolute allelic effects in the brain than in the liver (mean absolute allelic effect, brain versus liver = 0.215 versus 0.075; paired *t*-test: *t*(7) = 6.15, P = 4.7 × 10⁻⁴; Figure 8b), whereas the corresponding difference was not statistically significant for antagonistically pleiotropic loci (mean absolute allelic effect, brain versus liver = 0.138 versus 0.089; paired *t*-test: *t*(8) = 1.28, P = 0.24; Figure 8a).

**Fig. 8.**
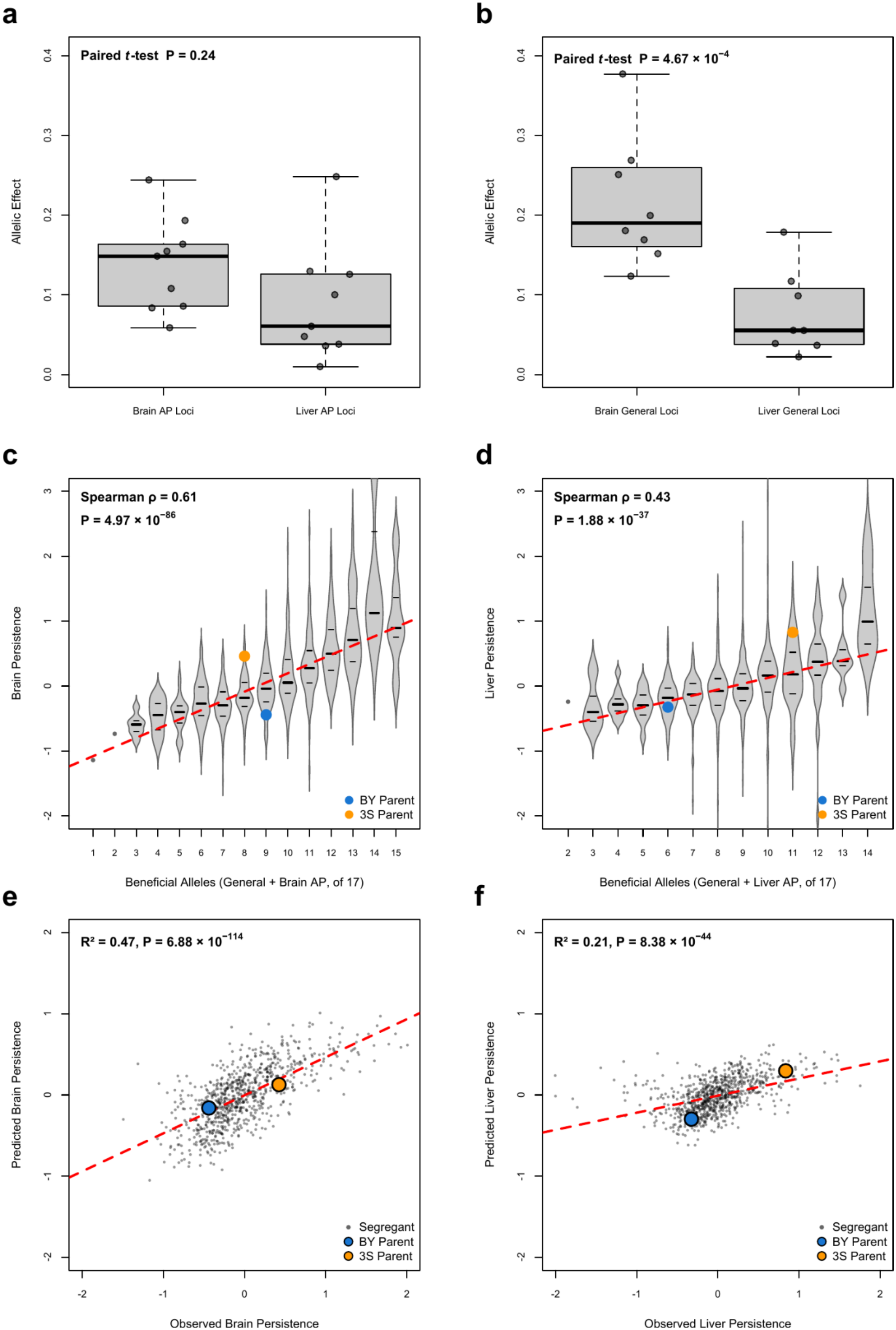
Allelic effects and additive-model predictions for the 17 aggregate loci. **a)** Magnitude of allelic effect in brain and liver for antagonistically pleiotropic loci (n = 9). **b)** The same for general loci (n = 8). Statistics, paired *t*-tests comparing absolute allelic effects between organs. Points are individual loci; boxes show the median and interquartile range, and whiskers extend to the most extreme values within 1.5 times the interquartile range. **c)** Brain persistence (mean across retained samples) against the number of brain-beneficial alleles carried across the 17 loci (beneficial alleles at general loci plus brain-beneficial alleles at antagonistically pleiotropic loci). **d)** Liver persistence against the equivalent liver count. Violins show the distribution at each count, with median and quartile bars. Black text, Spearman ρ and P value across the 816 genotyped segregants. Dashed red line, least-squares fit; blue and orange, BY and 3S parents. **e)** Predicted (y-axis) against observed (x-axis) brain persistence for each segregant under an additive model of the 17 loci. Observed persistence is mean fitted persistence from the fixed-effects model, averaged across host genotypes. The models were fit to the 816 genotyped segregants, so segregant predictions are in-sample fitted values; the parents were not included in the fit and were predicted from their genotypes. Black text, R^2^ and P value of the least-squares regression of predicted on observed persistence. **f)** The same for liver. Dashed red line, least-squares fit; blue and orange, BY and 3S parents. Alt text: Six panels in three rows. Top: box plots of allelic effect size in brain and liver for antagonistically pleiotropic loci (left, difference not statistically significant) and general loci (right, larger in brain). Middle: violin plots of brain (left) and liver (right) persistence against the number of beneficial alleles a segregant carries, with persistence rising with allele count and the parents marked in blue (BY) and orange (3S). Bottom: scatter plots of predicted against observed persistence under an additive model of 17 loci for brain (left) and liver (right), with a stronger relationship in brain.

Beneficial-allele counts at both locus classes were positively associated with persistence in each organ. In the brain, the number of beneficial general alleles carried by a segregant had a larger estimated correlation with persistence (Spearman ρ = 0.527, P = 1.3 × 10⁻⁵⁹) than the number of beneficial antagonistically pleiotropic alleles (ρ = 0.396, P = 5.9 × 10⁻³^2^). In the liver, the pattern was reversed: the number of beneficial antagonistically pleiotropic alleles carried by a segregant had a larger estimated correlation with persistence (ρ = 0.325, P = 1.8 × 10⁻^2^¹) than the number of beneficial general alleles (ρ = 0.237, P = 7.1 × 10⁻¹^2^; Supplementary Figure 3). The total number of organ-beneficial alleles carried by a segregant was significantly associated with persistence in both organs (brain: ρ = 0.615, P = 5.0 × 10⁻⁸⁶; liver: ρ = 0.427, P = 1.9 × 10⁻³⁷; Figure 8c,d). Together, these results are consistent with cumulative contributions from both locus classes to persistence in both organs, with general loci having larger average effects in the brain than in the liver.

### Identified loci partly explain the 3S advantage in brain and liver

To test whether the 17 loci could help explain the persistence advantage of 3S over BY, we fit an additive model for each organ in which the mean fitted persistence of the 816 genotyped segregants, averaged across host genotypes, was regressed on their genotypes at the peak markers of the 17 loci. These models explained 47% of the variance in mean fitted brain persistence (95% CI: 0.43–0.53) and 21% in mean fitted liver persistence (95% CI: 0.17–0.29; Figure 8e,f). Neither parent was used to fit the models. We then used the models to predict the persistence of BY and 3S from their genotypes at the 17 loci. The models predicted higher persistence for 3S than for BY in both organs, with a predicted difference of 0.29 in the brain (95% CI: 0.09–0.48) and 0.60 in the liver (95% CI: 0.42–0.79). 3S was predicted to exceed BY in 99.9% and 100% of 1,000 bootstrap replicates in brain and liver, respectively, with the 17 loci held fixed during resampling. These predicted differences were smaller than the observed differences in mean fitted persistence from the fixed-effects model (0.86 in the brain and 1.16 in the liver), so the additive models captured the direction of the 3S advantage but only about one-third of its magnitude in the brain and one-half in the liver.

The predicted advantage of 3S was consistent with differences in the number and estimated effects of beneficial alleles carried by the parents. Although BY carried more brain-beneficial alleles than 3S (9 versus 8), 3S showed greater brain persistence. At general loci, which had comparatively large average effects in the brain, 3S carried five beneficial alleles compared with three in BY. The additional brain-beneficial alleles carried by BY were at antagonistically pleiotropic loci, which had smaller average effects in the brain. In contrast, the liver advantage of 3S was consistent with its nearly twofold excess of liver-beneficial alleles relative to BY (11 versus 6; Supplementary Table 11). In the liver, mean absolute effect sizes were more similar between general and antagonistically pleiotropic loci (0.075 versus 0.089). 3S carried more beneficial alleles than BY at both general loci (5 versus 3) and antagonistically pleiotropic loci (6 versus 3), consistent with contributions from both classes to its larger persistence advantage in the liver (Figure 8c,d).

## Discussion

Our results support a largely shared genetic architecture of relative *Saccharomyces cerevisiae* persistence across the seven mouse host genotypes examined. Yeast genotype accounted for substantially more phenotypic variation than yeast genotype-by-host genotype interactions in both the brain and liver. Although host-dependent effects were statistically detectable, they explained a very small fraction of variation in the analyzed samples. Thus, under the conditions tested, host genetic differences did not substantially affect the overall genetic architecture of persistence. However, our assay measures relative persistence among yeast genotypes within hosts, not absolute fungal burden. Our results therefore cannot address whether absolute fungal burden or overall susceptibility to infection differs across mouse genotypes.

The linkage-mapping results further support the conclusion that genetic effects on relative persistence in the BY × 3S cross are largely shared across host genotypes. Loci detected across more host genotypes generally explained more within-host variance in the corresponding mapping phenotype, whereas each locus detected in only one host genotype explained less than 3% of that variance. Mapping mean persistence across hosts also recovered many genomic regions identified in our previous study using C57BL/6J mice, supporting the reproducibility of these associations within the BY × 3S cross (Mullis et al. 2022). Nevertheless, detection in only one host should not be equated with a genuinely host-specific allelic effect (Gelman and Stern 2006). The number of samples that passed quality control checks differed among host–organ combinations, and small-effect loci may fall below the detection threshold in some hosts. Differences in locus detection across hosts could therefore reflect small allelic effects and limited statistical power, rather than necessarily indicating host dependence.

Although genetic effects on persistence were largely shared across host genotypes, they differed between the brain and liver. As in our previous study, we identified both general loci, at which the same parental allele was associated with greater persistence across organs, and loci with opposing effects between organs, which we classified as antagonistically pleiotropic (Mullis et al. 2022). These patterns are consistent with genetic trade-offs between persistence in different parts of the mammalian body. The two classes also differed in their contributions to persistence. On average, general loci had larger effect sizes than antagonistically pleiotropic loci in the brain; however, this was not true in the liver. Together, these findings indicate that fungal genetic effects can be broadly shared across host genotypes yet differ in direction and magnitude between organs.

The parental comparisons and transgressive segregation further illustrate the polygenic basis of persistence. Although the clinical parent, 3S, had higher mean relative persistence than the laboratory parent, BY, in every host–organ combination, both parents carried beneficial alleles, and segregant phenotypes extended beyond the parental values in both organs. This pattern is consistent with recombination producing combinations of parental alleles that result in more extreme phenotypes (Brem and Kruglyak 2005). Among the mapped loci, 3S carried fewer brain-beneficial alleles overall but more beneficial alleles at general loci, which had comparatively large estimated effects in the brain. In the liver, 3S carried more beneficial alleles from both classes. These observations highlight the importance of allelic effect sizes and organ dependence, rather than beneficial-allele counts alone. Additive models fitted to the segregants predicted the direction, but not the full magnitude, of the parental difference in both organs. Together, these results suggest that 3S’s beneficial alleles at general loci may offset the brain-persistence costs of liver-beneficial alleles, helping explain its persistence advantage over BY in both organs despite antagonistic pleiotropy.

Future work could connect the mapped regions to the biological processes underlying persistence and test how broadly their effects generalize. Resolving the causal genes and variants would help determine whether general and antagonistically pleiotropic loci affect distinct processes, and whether opposing effects across organs arise from individual variants or from linked variants with different effects (Jiang and Zeng 1995). The current mapping intervals do not distinguish these possibilities. Extending the analysis to additional yeast genetic backgrounds and other mammalian species, potentially including rats or rabbits, would test whether these findings generalize beyond the BY × 3S cross and mouse hosts. In addition, pairing relative persistence measurements with measures of absolute fungal burden and host response could clarify how the genetic effects detected here relate to infection outcomes. Together, these studies could clarify how shared genetic effects and organ-specific trade-offs shape fungal persistence and its consequences for mammalian hosts.

## Data availability

Raw barcode sequencing reads are available from the NCBI Sequence Read Archive under BioProject PRJNA1549244. Segregant genotypes and barcode assignments are from Mullis et al. (2022). Processed barcode counts, per-sample persistence values, and linkage-mapping results are provided as Supplementary Tables 1–12 [and Supplementary Data]. Barcode counts, processed data, analysis code and supplementary material are available at the GSA figshare portal.

## Acknowledgments

We thank Alexandra Christensen and Ilan Goldstein for help with plating yeast extracted from mouse organ samples. We also thank Joseph Hale for assistance with the barcode sequencing pipeline. We thank Alexandra Christensen, Jinye Liang and Zachary Krieger for providing feedback on drafts of the manuscript. Figure 1 was created in Biorender (https://BioRender.com).

## References

Almeida F, Rodrigues ML, Coelho C. 2019. The Still Underestimated Problem of Fungal Diseases Worldwide. Front Microbiol. 10 [accessed 2026 Nov 2]. https://www.frontiersin.org/journals/microbiology/articles/10.3389/fmicb.2019.00214/full. 10.3389/fmicb.2019.00214

Ashman RB, Fulurija A, Papadimitriou JM. 1996. Strain-dependent differences in host response to Candida albicans infection in mice are related to organ susceptibility and infectious load. Infect Immun. 64(5):1866–1869. 10.1128/iai.64.5.1866-1869.1996

Bates D, Mächler M, Bolker B, Walker S. 2015. Fitting Linear Mixed-Effects Models Using lme4. J Stat Softw. 67:1–48. 10.18637/jss.v067.i01

Brem RB, Kruglyak L. 2005. The landscape of genetic complexity across 5,700 gene expression traits in yeast. Proc Natl Acad Sci U S A. 102(5):1572–1577. 10.1073/pnas.0408709102

Brown GD et al. 2012. Hidden Killers: Human Fungal Infections. Sci Transl Med. 4(165):165rv13–165rv13. 10.1126/scitranslmed.3004404

Byron JK et al. 1995. Pathogenicity of Saccharomyces cerevisiae in complement factor five-deficient mice. Infect Immun. 63(2):478–485. 10.1128/iai.63.2.478-485.1995

Churchill GA et al. 2004. The Collaborative Cross, a community resource for the genetic analysis of complex traits. Nat Genet. 36(11):1133–1137. 10.1038/ng1104-1133

Churchill GA, Doerge RW. 1994. Empirical threshold values for quantitative trait mapping. Genetics. 138(3):963–971. 10.1093/genetics/138.3.963

Churchill GA, Gatti DM, Munger SC, Svenson KL. 2012. The Diversity Outbred mouse population. Mamm Genome Off J Int Mamm Genome Soc. 23(9–10):713–718. 10.1007/s00335-012-9414-2

Ciurea CN et al. 2020. Candida and Candidiasis—Opportunism Versus Pathogenicity: A Review of the Virulence Traits. Microorganisms. 8(6):857. 10.3390/microorganisms8060857

Clemons KV, McCusker JH, Davis RW, Stevens DA. 1994. Comparative Pathogenesis of Clinical and Nonclinical Isolates of Saccharomyces cerevisiae. J Infect Dis. 169(4):859–867. 10.1093/infdis/169.4.859

Collaborative Cross Consortium. 2012. The Genome Architecture of the Collaborative Cross Mouse Genetic Reference Population. Genetics. 190(2):389–401. 10.1534/genetics.111.132639

Gelman A, Stern H. 2006. The Difference Between “Significant” and “Not Significant” is not Itself Statistically Significant. Am Stat. 60(4):328–331. 10.1198/000313006X152649

Goldstein AL, McCusker JH. 2001. Development of Saccharomyces cerevisiae as a Model Pathogen: A System for the Genetic Identification of Gene Products Required for Survival in the Mammalian Host Environment. Genetics. 159(2):499–513. 10.1093/genetics/159.2.499

Jiang C, Zeng ZB. 1995. Multiple Trait Analysis of Genetic Mapping for Quantitative Trait Loci. Genetics. 140(3):1111–1127. 10.1093/genetics/140.3.1111

Kivioja T et al. 2012. Counting absolute numbers of molecules using unique molecular identifiers. Nat Methods. 9(1):72–74. 10.1038/nmeth.1778

Köhler JR, Casadevall A, Perfect J. 2015. The Spectrum of Fungi That Infects Humans. Cold Spring Harb Perspect Med. 5(1):a019273. 10.1101/cshperspect.a019273

Levy SF et al. 2015. Quantitative evolutionary dynamics using high-resolution lineage tracking. Nature. 519(7542):181–186. 10.1038/nature14279

Liti G et al. 2009. Population genomics of domestic and wild yeasts. Nature. 458(7236):337–341. 10.1038/nature07743

Liu X et al. 2019. iSeq 2.0: A Modular and Interchangeable Toolkit for Interaction Screening in Yeast. Cell Syst. 8(4):338–344.e8. 10.1016/j.cels.2019.03.005

Marakalala MJ et al. 2013. Differential Adaptation of Candida albicans In Vivo Modulates Immune Recognition by Dectin-1. PLOS Pathog. 9(4):e1003315. 10.1371/journal.ppat.1003315

Matsui T et al. 2022. The interplay of additivity, dominance, and epistasis on fitness in a diploid yeast cross. Nat Commun. 13(1):1463. 10.1038/s41467-022-29111-z

McCusker JH. 2006. Saccharomyces cerevisiae: an Emerging and Model Pathogenic Fungus. In: Molecular Principles of Fungal Pathogenesis. John Wiley & Sons, Ltd; p 245–259 [accessed 2026 Nov 2]. https://onlinelibrary.wiley.com/doi/abs/10.1128/9781555815776.ch18. 10.1128/9781555815776.ch18

McCusker JH, Clemons KV, Stevens DA, Davis RW. 1994. Genetic characterization of pathogenic Saccharomyces cerevisiae isolates. Genetics. 136(4):1261–1269. 10.1093/genetics/136.4.1261

Morard M et al. 2023. Comparative genomics of infective Saccharomyces cerevisiae strains reveals their food origin. Sci Rep. 13(1):10435. 10.1038/s41598-023-36857-z

Mullis MN et al. 2022. Complex genetics cause and constrain fungal persistence in different parts of the mammalian body. Genetics. 222(3):iyac138. 10.1093/genetics/iyac138

Nivoix Y, Ledoux M-P, Herbrecht R. 2020. Antifungal Therapy: New and Evolving Therapies. Semin Respir Crit Care Med. 41(1):158–174. 10.1055/s-0039-3400291

Radovanovic I, Mullick A, Gros P. 2011. Genetic Control of Susceptibility to Infection with Candida albicans in Mice. PLOS ONE. 6(4):e18957. 10.1371/journal.pone.0018957

Scorzoni L et al. 2017. Antifungal Therapy: New Advances in the Understanding and Treatment of Mycosis. Front Microbiol. 8 [accessed 2026 Nov 2]. https://www.frontiersin.org/journals/microbiology/articles/10.3389/fmicb.2017.00036/full. 10.3389/fmicb.2017.00036

Strope PK et al. 2015. The 100-genomes strains, an S. cerevisiae resource that illuminates its natural phenotypic and genotypic variation and emergence as an opportunistic pathogen. Genome Res. 25(5):762–774. 10.1101/gr.185538.114

Tuite A et al. 2005. Genetic control of susceptibility to Candida albicans in susceptible A/J and resistant C57BL/6J mice. Genes Immun. 6(8):672–682. 10.1038/sj.gene.6364254

van de Veerdonk FL et al. 2025. Aspergillus fumigatus biology, immunopathogenicity and drug resistance. Nat Rev Microbiol. 23(10):652–666. 10.1038/s41579-025-01180-z

Zhao L, Liu Z, Levy SF, Wu S. 2018. Bartender: a fast and accurate clustering algorithm to count barcode reads. Bioinformatics. 34(5):739–747. 10.1093/bioinformatics/btx655

